# Redirecting resistance evolution in *BRAF^V600^* melanoma by inhibition of the peroxiredoxin-thioredoxin system

**DOI:** 10.1101/2025.03.10.641595

**Authors:** Stefanie Egetemaier, Heike Chauvistré, Renáta Váraljai, Yichao Hua, Smiths S. Lueong, Samira Makhzami, Nalini Srinivas, Jan Forster, Vivien Ullrich, Simone Stupia, Valéria Schroeder, Sarah Scharfenberg, Anna Hoewner, Batool Shannan, Jens Siveke, Maria Francesca Baietti, Eleonora Leucci, Jean-Christophe Marine, Annette Paschen, Bjoern Scheffler, Daniel R. Engel, Lisa M. Becker, Kathrin Thedieck, Felix Nensa, Johannes Koester, Barbara T. Grünwald, Simon Poepsel, Sabrina Ninck, Farnusch Kaschani, Dirk Schadendorf, Juergen C. Becker, Alpaslan Tasdogan, Florian Rambow, Alexander Roesch

**Author notes:** These authors contributed equally (Stefanie Egetemaier, Heike Chauvistré). These authors contributed equally (Florian Rambow, Alexander Roesch).

## Abstract

Drug-tolerant persister cells (DTPs) exhibit remarkable cell state heterogeneity and phenotypic evolvability. However, the central question of how DTPs epigenetically coordinate their metabolic flexibility to adapt early to therapeutic stress remains unanswered. We have recently shown that the histone demethylase KDM5B, which is intrinsically expressed in differentiated melanoma DTPs, reprograms the metabolic cell landscape. However, the exact mechanism by which KDM5B affects underlying metabolic enzymes, and whether this reveals new druggable vulnerabilities, remain unknown. By transcriptional and epigenetic profiling of *BRAFV600* melanoma cells following KDM5B gene silencing, we discovered a direct molecular axis between the epigenetic regulator KDM5B and the PRDX/TXN ROS detoxification system. This metabolic axis is regulated independently of KDM5B’s demethylase activity. Furthermore, RNAi approaches and the pharmacological inhibition of the PRDX/TXN system led to ROS-induced cell death in differentiated melanoma DTPs and a delay of resistance development to MAPK inhibition. This process was independent of lipid-ROS-driven ferroptosis. Additionally, single-cell transcriptome analyses from pre-clinical melanoma models under continuous MAPK inhibitory treatment demonstrated altered cellular differentiation dynamics, with a reduction in the early evolution into the mesenchymal DTP state under concomitant PRDX inhibition. Interestingly, the degree of melanoma cell state differentiation at the onset of treatment was a major determinant for the transition towards the neural crest-like DTP state. Our study identified a high degree of epigenetic-metabolic connectivity and flexibility within the melanoma DTP pool and urges caution with single redox pathway-targeted strategies for tumor elimination in the future. Prospectively, our results point towards a new resistance targeting strategy for *BRAFV600* melanoma patients based on pharmacological re-direction of the evolution of melanoma cell states already at therapy onset.

**Highlights:**

- Transcriptional and epigenetic profiling identified KDM5B as a regulator of the PRDX/TXN-ROS detoxification system
- PRDX inhibition increases the vulnerability of KDM5B^high^ DTPs to ROS, independent of ferroptosis
- PRDX inhibition delays resistance to MAPK inhibition in *BRAFV600* melanoma cells
- Longitudinal single-cell transcriptome analysis reveals that PRDX inhibition re-directs early DTP evolution
- PRDX/TXN gene expression is predictive for melanoma patient survival

## Introduction

Tumor cells face multiple challenges to escape from hostile conditions^1^. While the majority of tumor cells is susceptible to environmental or therapeutic stress, some subpopulations of tumor cells persist due to phenotypic (non-genetic) traits that ensure their long-term survival and promote constant tumor evolution^2–7^. In particular, melanoma drug-tolerant persister cells (DTPs) exhibit remarkable cell state heterogeneity and metabolic adaptability when exposed to exogenous stress, including chemotherapy, ionizing radiation, or, more specific to melanoma cell function, mitogen-activated protein kinase (MAPK)-targeted treatment (BRAF/MEK inhibition)^2,8–10^. However, the central question remains unanswered as to how melanoma DTPs epigenetically coordinate metabolic flexibility to dynamically adapt to such manifold toxic stressors.

On the cellular differentiation level, melanoma DTPs represent a heterogeneous cell pool covering a broad range of mesenchymal, neural crest-like, and differentiated cell identities that can dynamically transition into each other^2,11–13^. We and others recently showed that the dynamics in therapy persistence of melanoma cells are possible through the epigenetic control of cellular energy homeostasis^8,14–16^. For example, we demonstrated that a subpopulation of melanoma DTPs can rapidly balance their oxidative energy supply by upregulating the chromatin regulator KDM5B^8^. KDM5B controls multiple transcriptional programs by histone H3 lysine 4 (H3K4) demethylation, but has recently been recognized also for its demethylase-independent functions^17,18^.

We have previously found that KDM5B drives the switch from glycolytic to oxidative phosphorylated (OXPHOS) ATP as preferred energy source in differentiated melanoma DTP cells^8,13,16^. However, pharmacological inhibition of KDM5s only showed limited efficacy in terms of tumor growth inhibition^16^. Also, the exploitation of OXPHOS dependence as a therapeutic vulnerability in preclinical and clinical melanoma studies has not been very successful, as the therapeutic index of pharmacological OXPHOS inhibitors is very narrow, with high dose-limiting toxicities and dynamic metabolic adaptations of surviving melanoma cells at the concentrations used^19,20^. Similarly, blocking of mitochondrial translation with tetracyclines, although very efficient at eradicating dedifferentiated DTPs, failed to efficiently eliminate differentiated cells^21^.

Another DTP cell vulnerability that has been identified across different cancer entities is the cellular detoxification of reactive oxygen species (ROS)^22–24^. Cellular ROS levels can be elevated in cancer cells due to multiple mechanisms including ROS-generating nicotinamide adenine dinucleotide phosphate (NADPH)-dependent oxidases (NOXs), through beta-oxidation of fatty acids in the peroxisome, but mainly as a result of mitochondrial leakage of electrons due to enhanced OXPHOS^25–28^. Accordingly, GPX4-mediated reduction of reactive (lipid) peroxides (lipid ROS) to stable, non-toxic lipid alcohols has been shown to promote DTP cell survival in breast, ovarian, lung cancer, and also melanoma^29,30^. Pharmacological inhibition of GPX4 can induce ferroptosis, a cell death modality driven by toxic lipid ROS levels^31^. Ferroptosis-inducing compounds such as the GPX4 inhibitor RSL3 show particularly high efficacy in dedifferentiated, mesenchymal DTPs^12^; however, poor pharmacokinetics and a lack of GPX4-selectivity currently constrain the clinical application of GPX4 inhibitors^32^.

In this study, we identified a novel molecular axis between the epigenetic regulator KDM5B and the peroxiredoxin (PRDX)-thioredoxin (TXN) ROS detoxification system as an alternative to GPX4-driven redox balancing. Furthermore, the PRDX/TXN system had a direct effect on the cell differentiation state of melanoma DTPs and may thus represent a novel target downstream of KDM5B. Indeed, pharmacological inhibition of PRDX increased the vulnerability of melanoma DTPs to ROS and attenuated the transcriptional tumor evolution towards dedifferentiated DTP states under prolonged MAPK inhibition, thereby overcoming the known limitations of direct KDM inhibition and of the low bioavailability of current GPX4 inhibitors. Our results emphasize not only a high degree of epigenetic-metabolic connectivity and flexibility within the melanoma DTP pool but also urge caution with single redox pathway-targeted strategies for tumor elimination in the future.

## Results

### KDM5B-dependent regulation of metabolic gene signatures

Using liquid chromatography-high resolution mass spectrometry profiling of cellular metabolites, KDM5B was recently suggested to reprogram melanoma cells towards increased oxidative and redox metabolism^16^. However, the exact mechanism by which KDM5B affects the underlying metabolic enzymes and what this means for the biological functionality of cancer cells remain unknown. To identify a potential KDM5B-dependent epigenetic-metabolic axis and define the involved KDM5B-dependent target enzymes, we performed transcriptional profiling of melanoma cells after gene silencing. Bulk RNA sequencing (RNA-seq) was performed following doxycycline induced KDM5B knockdown in WM3734 melanoma cells (Figure S1A). Interestingly, Gene Set Enrichment Analysis (GSEA) of differentially regulated transcriptional signatures after KDM5B knockdown found the downregulation of a number of metabolic pathways (Figure 1A blue bars, 1B, S1B). In particular, genes from “HALLMARK OXIDATIVE PHOSPHORYLATION” and “REACTOME DETOXIFICATION OF REACTIVE OXYGEN SPECIES” were downregulated after knockdown of KDM5B. Subsequent re-analysis of publicly available single cell transcriptomes from 19 patients confirmed a correlation of high KDM5B expression with OXPHOS and ROS detoxification genes in the melanoma tissue context (data from 1252 melanoma cells^33^) (Figure 1C, S1C). Notably, within the top significantly regulated ROS detoxification genes, four genes, PRDX1, PRDX3, PRDX5, and TXN, belonged to the same redox pathway, i.e. the peroxiredoxin-thioredoxin (PRDX/TXN) system^34^. Since the PRDX/TXN system has not been systematically studied in the context of melanoma, we conducted a re-analysis of the TCGA_SKCM dataset to investigate its general relevance for patient survival. Interestingly, most of the individual PRDX/TXN genes were associated with an increased risk for death and also the combined gene expression signature across all PRDX/TXN genes was significantly associated with reduced overall survival (Figure 1D).

**Figure 1:**
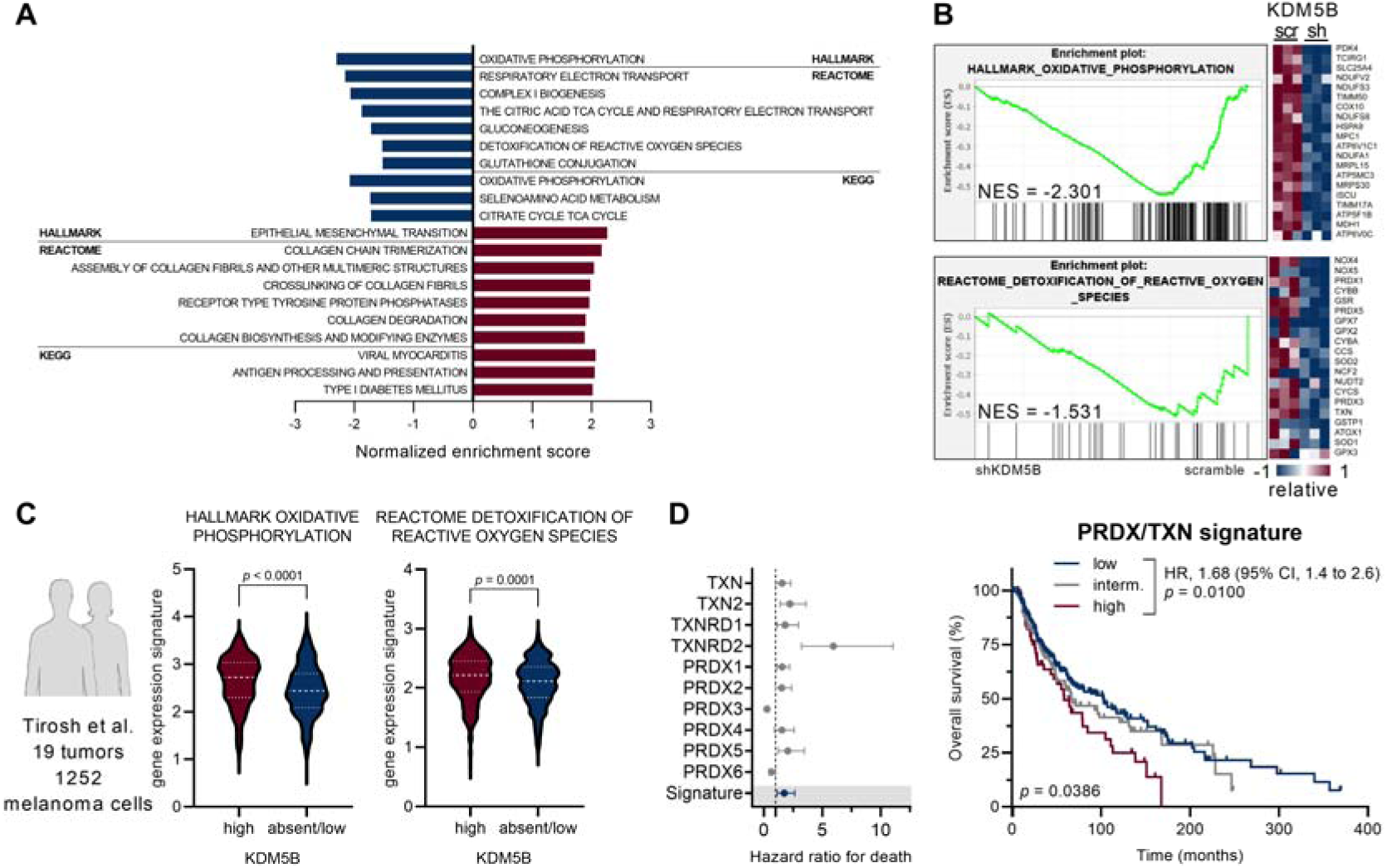
KDM5B-dependent regulation of oxidative metabolism gene signatures. **A** Bar graph showing significant metabolic gene sets from Hallmark, Reactome, and KEGG database with nominal *p*-value <0.05 and normalized enrichment score <-1.5 and >1.5. Downregulated gene sets in Dox-Tet3G-induced shKDM5B knockdown vs. scramble (blue bars), upregulated gene sets (red bars) **B** Representative GSEA enrichment plots for the indicated gene sets and heatmap representation of the top 20 ranked genes. Gene expression levels are displayed on a relative color scale from low (blue) to high (red). **C** Violin expression plots of the indicated gene sets relative to KDM5B in single-cell RNA-sequenced human melanoma cells^33^. Significance was tested by Mann Whitney test. **D** Overall survival of cutaneous melanoma patients (TCGA_SKCM cohort) in relation to peroxiredoxin-thioredoxin (PRDX/TXN) gene expression. Cut-point optimization for low, intermediate (interm.), and high expression was calculated for each gene with X-tile. Forest plot showing the hazard ratio for death and 95% confidence intervals for individual PRDX/TXN genes and PRDX/TXN signature (left panel). Kaplan-Meier analysis of PRDX/TXN signature. Significance was tested by Log-rank (Mantel-Cox) test. Overall significance *p* = 0.0386, low vs. high *p* = 0.0100 (right panel). Abbreviation: NES: normalized enrichment score. See also Figure S1.

### KDM5B-dependent regulation of PRDX/TXN genes

Having identified the PRDX/TXN system as potential downstream target of KDM5B, we proceeded to assess how KDM5B regulates PRDX/TXN genes. As a known histone H3 lysine 4 demethylase, KDM5B removes di- and tri-methyl groups (H3K4me2/3)^18^. Thus, H3K4me3-chromatin immunoprecipitation sequencing (H3K4me3-ChIP-seq) was performed in the same doxycycline inducible KDM5B knockdown cells. However, unexpectedly, we could neither find significant differences in peak signal intensities nor peak calling between the scramble control and KDM5B knockdown cells for PRDX/TXN genes (Figure S2A). To confirm that the selected H3K4me3-ChIP-seq approach is in principle capable of detecting histone methylation differences depending on KDM5B levels, we checked gene regions with known functions in cell state dedifferentiation in melanoma^12^. According to our previous observations, KDM5B promotes melanocytic differentiation programs while repressing dedifferentiation gene expression^13^. Accordingly, H3K4me3-ChIP-seq revealed an increase in peak signal intensities and peak calling in KDM5B knockdown cells at expected exemplary gene regions like AXL, SPOCK1, and PXDN (Figure S2B). Consequently, PRDX/TXN genes might not be epigenetically regulated by KDM5B’s demethylase activity. Nevertheless, based on KDM5B-ChIP-seq in melanoma cells and *in silico* analysis of ChIP-seq data from the ChIP-Atlas^35^, KDM5B binding sites were present at the respective promoter regions of several PRDX/TXN genes including PRDX2, PRDX3, PRDX5, TXN2, and TXNRD2 (Figure 2A). To validate these findings, chromatin immunoprecipitation followed by quantitative PCR (ChIP-qPCR) was performed, confirming the presence of KDM5B binding sites at PRDX/TXN genes (Figure 2B), and suggesting a KDM5B demethylase-independent regulation of PRDX/TXN genes. In fact, KDM5B was previously shown to exhibit demethylase-independent functions as part of larger chromatin-associated protein complexes: for example, the *Drosophila* KDM5 ortholog regulates oxidative stress genes via interactions with HDAC4 and FOXO proteins^36^. To investigate a potentially similar interaction of human KDM5B with transcription factors (FOXO) or histone deacetylases (HDAC), we performed KDM5B-immunoprecipitation in nuclear lysates followed by mass spectrometric detection. Firstly, KDM5B enrichment was confirmed in the KDM5B immunoprecipitation eluate by western blotting (Figure S2C). Subsequent mass spectrometric analysis did not support a significant interaction with FOXO1 (Figure S2D), nor with any of the three other human FOXO paralogs (FOXO3, FOXO4, and FOXO6) while revealing a substantial binding to HDAC1 and HDAC2 (Figure S2D, S2E). *In silico* HDAC1- and HDAC2-ChIP-seq data from the ChIP-Atlas^35^ furthermore confirmed genomic HDAC binding sites overlapping with KDM5B binding at the same PRDX/TXN gene regions (Figure 2A). Additionally, multiplex siRNA gene knockdown of KDM5B, HDAC1, and HDAC2 downregulated most PRDX/TXN genes in different melanoma cell lines (Figure S2F), altogether, suggesting that KDM5B could be part of a functional active HDAC-containing chromatin complex.

**Figure 2:**
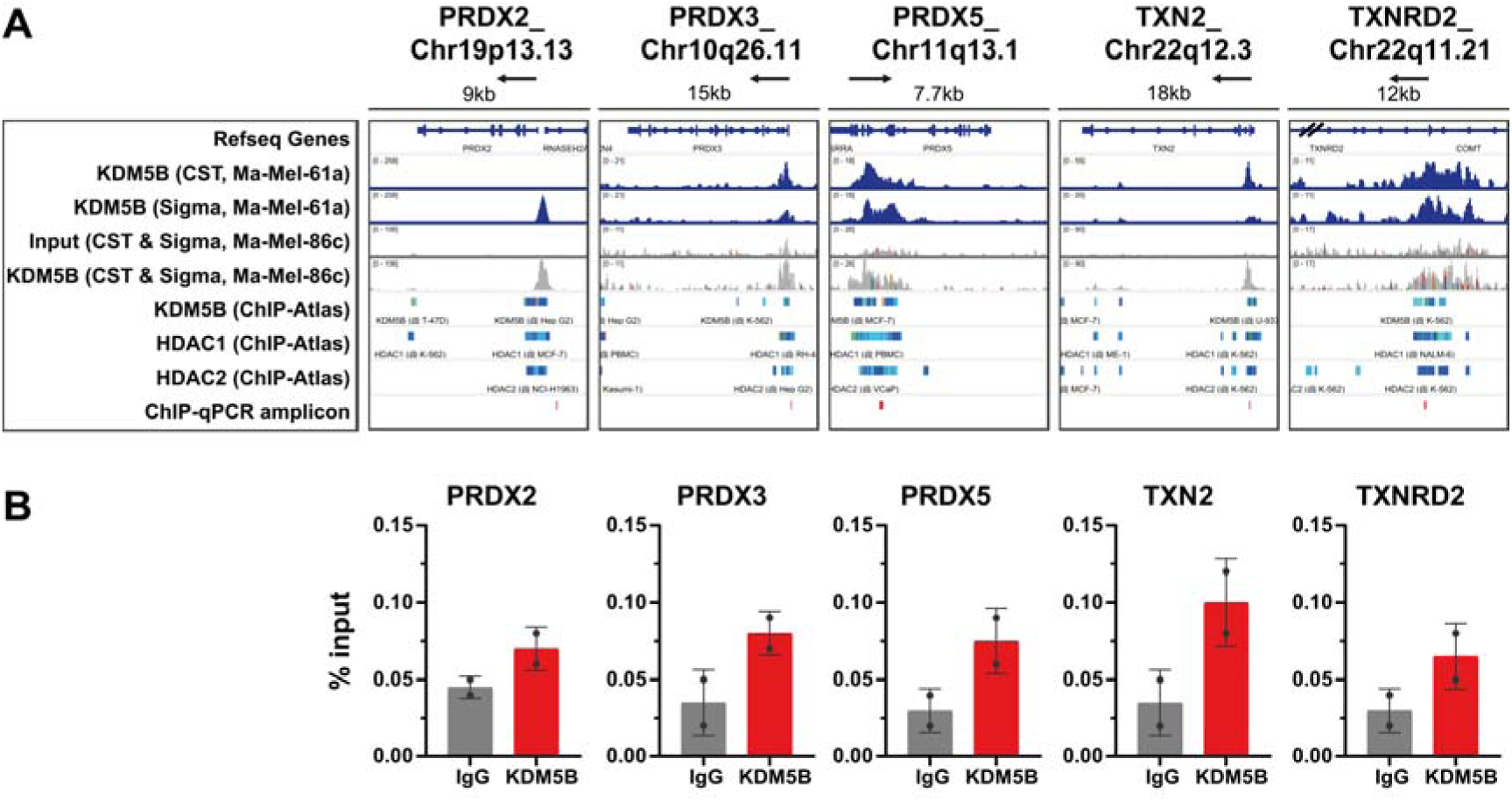
KDM5B-dependent regulation of PRDX/TXN genes. **A** KDM5B intensity peak signals from KDM5B-ChIP-seq data in naïve Ma-Mel-61a and Ma-Mel-86c cells (blue and grey peaks respectively), depiction of KDM5B, HDAC1, and HDAC2 binding from the ChIP-Atlas (color gradient bars)^35^, and amplicon regions of ChIP-qPCRs (red bars) at PRDX/TXN promoter regions. Data was visualized using the Integrative Genomics Viewer (IGV). Arrows indicate direction of transcription, // indicates truncated TXNRD2 gene. **B** KDM5B binding at promoter regions of indicated PRDX/TXN genes in ChIP-qPCR analysis in Ma-Mel-61a cells. Bar graphs represent % of input of IgG or KDM5B antibody. Data are represented as mean ± SD from 2 independent biological experiments. Abbreviations: CST: KDM5B antibody from Cell Signaling Technology, Sigma: KDM5B antibody from Sigma-Aldrich. See also Figure S2.

### Inhibition of PRDX/TXN gene functions limits melanoma cell viability

Previously, we have shown that expression of KDM5B is important for melanoma survival^3^. To investigate the potential impact of PRDX/TXN genes on melanoma cell viability, we established multiplex siRNA targeting of either all thioredoxins (TXN1-2), all thioredoxin reductases (TXNRD1-2), or all peroxiredoxins (PRDX1-6) (Figure S3A). siRNAs were transfected into WM3734 cells that harbored our previously validated KDM5B-promoter-EGFP-reporter construct (WM3734^KDM5Bprom-EGFP^ cells^3^). This allowed us to simultaneously monitor both KDM5B expression by EGFP levels and cell survival by 7-AAD staining via flow cytometry (Figure 3A). Baseline cell death after knockdown of all PRDX/TXN genes was moderate. However, when cells were additionally put under exogenous stress by a BRAF^V600E^-targeting drug (2.5 µM PLX4720), a strong additive effect was seen on cell killing leading to decreased levels of KDM5B-expressing (EGFP+) cells.

**Figure 3:**
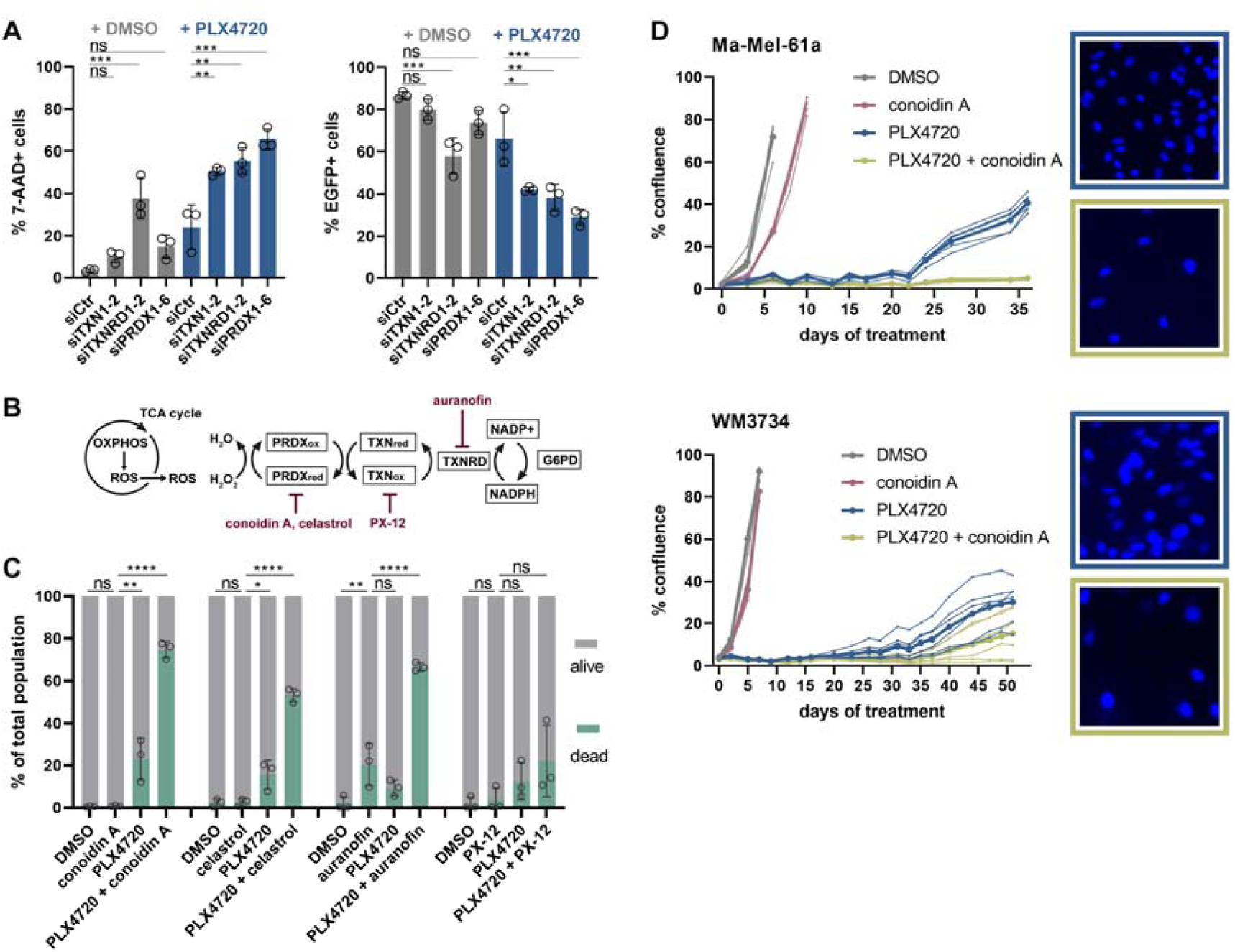
Gene knockdown and chemical inhibition of PRDX/TXN limits melanoma cell viability under exogenous stress. **A** Flow cytometric detection of 7-AAD and EGFP in WM3734^KDM5Bprom-EGFP^ cells. Cells were analyzed 96 hours after siRNA transfection of the indicated genes and 72 hours after DMSO or PLX4720 treatment. Left panel: 7-AAD quantitation. Right panel: EGFP quantitation. Significance was tested by ordinary one-way ANOVA with Dunnett’s multiple comparison test. Data are represented as mean ± SD from 3 independent biological experiments. **B** Schematic representation of the PRDX/TXN system and pharmacological inhibitors. **C** Quantitation of alive/dead WM3734^KDM5Bprom-EGFP^ cells by flow cytometric detection of 7-AAD after treatment with indicated drugs for 72 hours. Significance was tested by ordinary one-way ANOVA with Dunnett’s multiple comparison test. Data are represented as mean ± SD from 3 independent biological experiments. **D** Representative long-term *in vitro* repopulation assay with continuous treatment and confluence measurement in Ma-Mel-61a cells (top panel) and WM3734 cells (bottom panel). Confluence data are represented as individual 6 technical replicates and mean of the 6 technical replicates (thicker line). Representative pictures of Hoechst-stained nuclei at the end of the experiment on day 38 (Ma-Mel-61a) and day 51 (WM3734) (right panels). One representative experiment per cell line out of n=3. See also Figure S3.

To pharmacologically phenocopy the effects of genetic PRDX/TXN silencing, we assessed a variety of commercially available compounds known to block PRDX/TXN functions (Figure 3B). Auranofin (TXNRD inhibitor) is an FDA-approved drug for the treatment of rheumatoid arthritis^37^. PX-12 (TXN inhibitor) and celastrol (PRDX2 inhibitor) were or are currently investigated in clinical trials as therapies for advanced pancreatic cancer^38^ or as dietary supplements^39^, respectively, while conoidin A (PRDX1, PRDX2, and PRDX4 inhibitor) is used for treatment of pre-clinical prostate cancer models or patient-derived glioblastoma cells^40,41^. First, we determined IC_50_ values for all applied PRDX/TXN inhibitors to identify effective concentrations for subsequent experiments within a killing range comparable to the siRNA experiment (Figure S3B). In line with genetic silencing (Figure 3A), PRDX/TXN inhibition alone did not result in notable effects on cell viability for most inhibitors in the same reporter cell line, whereas combination with PLX4720 showed significant additive killing for most inhibitors (Figure 3C).

Performing the experiment with a clinically relevant combination of a BRAF^V600E^ inhibitor plus a MEK inhibitor (dabrafenib plus trametinib) confirmed this observation (Figure S3C). Due to the significantly higher toxicity associated with dual mitogen-activated protein kinase (MAPK) inhibition, we used lower compound concentrations in this experiment to receive measurable effects. Nevertheless, this resulted in a weaker effect size as compared to the combination with the BRAF^V600E^ inhibitor alone.

Given the pivotal role of PRDX within the PRDX/TXN system through the direct reduction of hydrogen peroxide^42^, we conducted individual gene knockdowns of the six PRDXs to explore PRDX isoform-specific roles in melanoma cell survival (Figure S3D). 7-AAD staining under PLX4720-induced exogenous stress revealed significant additive cell killing effects for PRDX2, PRDX4, PRDX6, and the most pronounced effect for PRDX3 (Figure S3E).

We next wondered if PRDX/TXN functions are also relevant for the previously reported self-renewal capacity of KDM5B-expressing melanoma cells^3^. We selected the PRDX inhibitor conoidin A as a tool compound for subsequent experiments. We set up a long-term assay in which we monitored cell survival and growth via confluence measurement under permanent BRAF^V600E^ inhibitor stress and/or conoidin A treatment in BRAF^V600E^-mutated Ma-Mel-61a or WM3734 cells (Figure 3D). In accordance with the short-term assays done earlier, conoidin A alone had no effect on melanoma cell proliferation, while the BRAF^V600E^ inhibitor initially caused strong cell elimination followed by resistance development and culture repopulation after 20-30 days (time varies depending on the cell line tested). Yet, combination of conoidin A and BRAF^V600E^ inhibitor considerably reduced culture repopulation. Hoechst staining of surviving cells’ nuclei at the end of the experiments confirmed conoidin A’s potential to reduce long-term growth in the combination with both BRAF inhibition and BRAF/MEK inhibition (Figure 3D, S3F).

In sum, our findings suggest that surviving melanoma DTPs sustain their viability by the PRDX/TXN system and inhibition of PRDX’s can significantly reduce melanoma persistence.

### PRDX inhibition increases ROS-dependent cell death independent of ferroptosis

Imbalanced high ROS levels have cell-damaging effects and limit cell viability^22^. We thus tested whether the observed increase in cell death after PRDX inhibition in BRAF^V600E^-targeted cells was related to increased ROS levels. We measured general ROS levels by CellROX™ green in PRDX/TXN inhibitor treated cells. In particular, the treatment with the PRDX inhibitors conoidin A and celastrol led to increased ROS in a time-dependent manner (Figure 4A). The antioxidant N-acetyl-L-cysteine (NAC), a ROS scavenger, rescued high ROS levels, even below the baseline ROS level of untreated cells. For PX-12, no major changes in ROS levels were observed, neither in single treatment nor upon NAC rescue. In cells treated with auranofin, a reduction of high ROS levels with NAC treatment was not seen. Also, when added to BRAF^V600E^ inhibition, conoidin A showed a time-dependent increase in ROS levels, which could be rescued by NAC at early treatment time points (Figure 4B).

**Figure 4:**
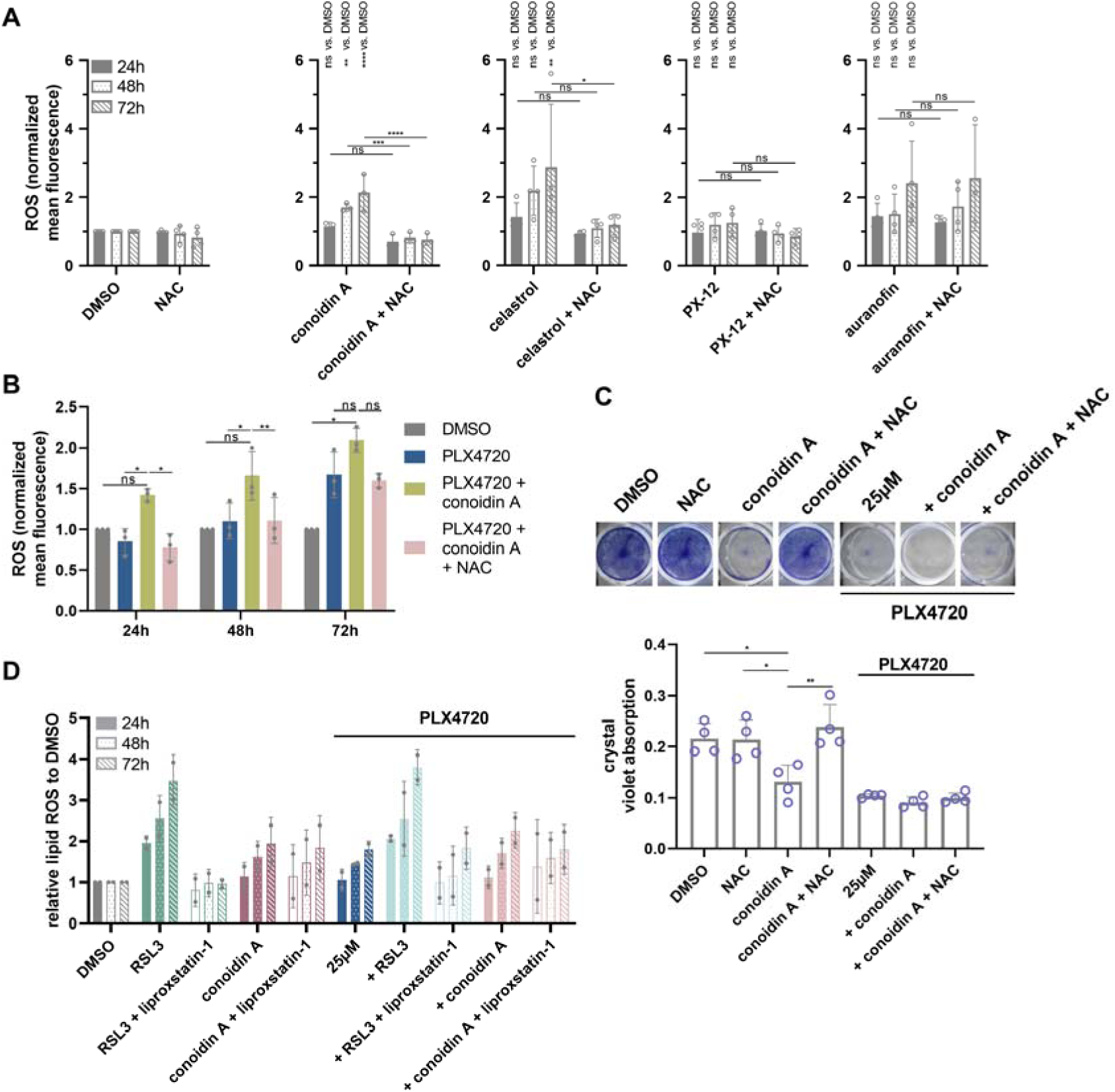
PRDX inhibition increases ROS-dependent cell death independent of ferroptosis. **A** ROS levels stained by CellROX™ green in Ma-Mel-86c cells after treatment for 24, 48, and 72 hours with indicated drugs. ROS levels were rescued with the ROS scavenger NAC. Mean fluorescence intensities were normalized to DMSO controls. Significance was tested by two-way ANOVA with Holm-Sidak’s multiple comparison test. Data are represented as mean ± SD from 3-4 independent biological experiments. **B** ROS levels stained by CellROX™ green in Ma-Mel-86c cells after treatment for 24, 48, and 72 hours with indicated drugs and NAC. Mean fluorescence intensities were normalized to DMSO controls. Significance was tested by two-way ANOVA with Tukey’s multiple comparison test. Data are represented as mean ± SD from 3 independent biological experiments. **C** Representative pictures (top) and crystal violet quantitation (bottom) of cell viability rescue experiment in Ma-Mel-86c cells treated for 72 hours with high conoidin A concentration (2.5 µM) and indicated drugs. Significance was tested by ordinary one-way ANOVA with Dunnett’s multiple comparison test. Data are represented as mean ± SD from 4 independent biological experiments. **D** Normalized levels of lipid ROS stained by C11 BODIPY 581/591 in Ma-Mel-86c cells after 24, 48, and 72 hours of treatment with indicated drugs. Data are represented as mean ± SD from 2 independent biological experiments. Abbreviation: NAC: N-acetyl-L-cysteine.

Next, we tested whether conoidin A-induced ROS have a direct effect on melanoma cell survival. As previously shown in Figure 3C, conoidin A is not toxic at the concentration used before (0.5 μM). To be able to measure cell death upon conoidin A treatment, we had to apply a high drug concentration (2.5 µM). Measurement of cell survival after 72 hours of conoidin A treatment led to a significant reduction in cell survival which could be fully restored when cells were additionally treated with NAC to rescue increased ROS. Similar effects of NAC were observed in conoidin A plus BRAF^V600E^ inhibitor-treated cells (Figure 4C).

Cancer cell persistence has been repeatedly reported to depend on ROS detoxification via GPX4 pointing towards ferroptosis as a common cancer-overarching vulnerability^43^. Since ferroptosis is caused by high levels of membrane lipid peroxidation, we applied C11 BODIPY 581/591 as fluorescence-based assay to detect lipid peroxides (Figure 4D). Melanoma cells treated with the ferroptosis inducer RSL3 showed a strong, time-dependent increase in lipid ROS, which could be rescued by the ferroptosis inhibitor liproxstatin-1. Interestingly, we observed a less than 2-fold increase in lipid ROS after treating cells with conoidin A. Additionally, liproxstatin-1 did not rescue the lipid ROS levels in conoidin A treated cells. Moreover, also the combination of conoidin A and PLX4720 only slightly increased C11 BODIPY 581/591 signals with again no rescue effect by liproxstatin-1. In sum, these findings suggest that conoidin A inhibition increases ROS-dependent cell death independent of ferroptosis.

### PRDX are required for melanoma cell state evolution

Melanoma cells can escape from therapeutic pressure stress by shifting between different cell states^11^. In particular, the dynamic evolution towards a dedifferentiated transcriptional phenotype is key for survival under MAPK inhibitory therapy^2^. To investigate how PRDX-related functions affect melanoma cell state shifts, we longitudinally exposed WM3734 cells to dabrafenib plus trametinib as clinically relevant drugs in combination with conoidin A. As expected based on previous reports^2^, expression of NGFR as a surrogate marker for the neural crest-like (NC-like) state significantly increased upon treatment with dabrafenib plus trametinib (Figure 5A). Upon additional treatment with conoidin A, NGFR expression also increased, although to a lesser extent than with dabrafenib plus trametinib treatment. In contrast, the melanocytic lineage marker MITF increased after conoidin A co-treatment (Figure S4A).

**Figure 5:**
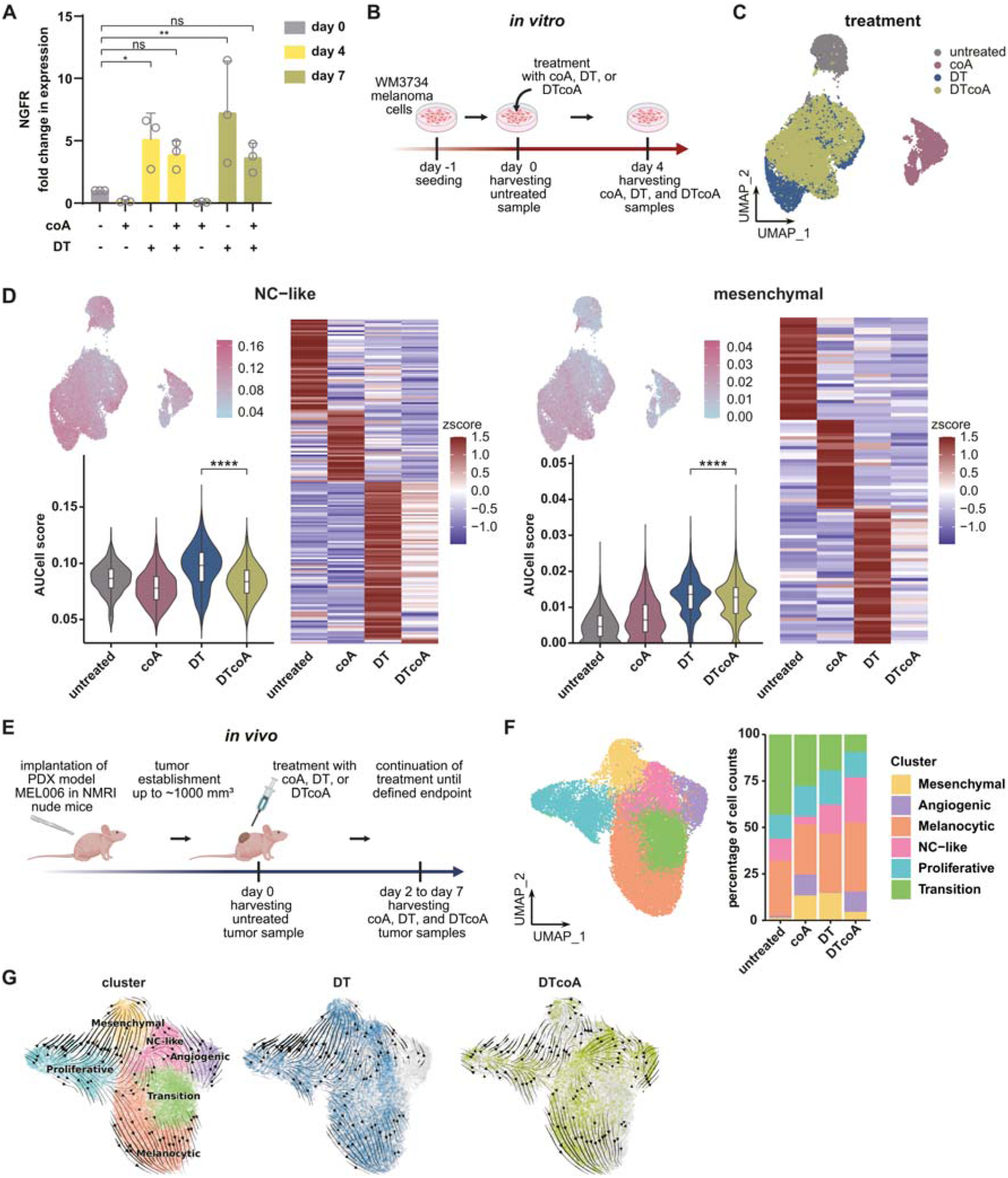
PRDXs are required for temporal cell state evolution. **A** Quantitation of NGFR gene expression by qPCR in WM3734 cells longitudinally treated with coA, DT, or DTcoA. Significance was tested by ordinary one-way ANOVA with Holm-Sidak’s multiple comparison test and a preselection of untreated, DT, and DTcoA samples. Data are represented as mean ± SD from 3 independent biological experiments. **B** Schematic representation of *in vitro* single-cell RNA sequencing (scRNA-seq) experiment. **C** UMAP generated from *in vitro* scRNA-seq data (B) color-coded by treatment. **D** UMAP, Violin-plus boxplot with median-quantile-min/max without outliers and population distribution, and heatmap generated from scRNA-seq data, displaying combined differentiation signatures (NC-like: left panel, mesenchymal: right panel) from ^12^ and ^44^. Statistics were assessed by Wilcoxon test with the adjusted *p*-value using Holm correction. **E** Schematic representation of *in vivo* single-nuclei RNA sequencing (snRNA-seq) experiment. **F** UMAP (left panel) showing cluster annotation of *in vivo* snRNA-seq experiment (E). Bar chart (right panel) showing the proportions of the six clusters. **G** RNA velocities (velocyto/scVelo) projected onto UMAP from snRNA-seq experiment. Cells are color-coded according to cluster (left panel) or treatment (middle and right panel). Arrows indicate the differentiation direction of cells in all four samples with cluster annotation (untreated, coA, DT, and DTcoA) (left panel) and in selected cells from DT (blue) or DTcoA (green) treatment samples (middle and right panel). Abbreviations: coA: conoidin A, DT: dabrafenib plus trametinib, DTcoA: dabrafenib plus trametinib plus conoidin A. See also Figure S4.

To elucidate the role of PRDX inhibition among the heterogeneous melanoma DTP states, single-cell RNA sequencing (scRNA-seq) was performed after short-term (4 days) treatment with conoidin A, dabrafenib plus trametinib, or their combination, alongside an untreated control (Figure 5B). UMAP-based dimension reduction and Louvain-clustering revealed “treatment type” as the predominant source of variance (Figure 5C). GSEA, utilizing gene signatures for NC-like and mesenchymal cell states sourced from two published studies^12,44^, demonstrated an induction of these signatures following dabrafenib plus trametinib treatment compared to the untreated condition (Figure 5D). Co-treatment with conoidin A led to a significant reduction in NC-like and mesenchymal signatures genes. SCENIC analysis^45^ of the underlying gene regulatory networks indicated increased activity of neuroectodermal and mesenchymal transcription factors (TFs) such as SOX6, SOX10, MEF2C, SOX4, JUN, JUND, and ETS1 following dabrafenib plus trametinib treatment, with a subsequent decrease upon conoidin A addition (Figure S4B), suggesting their involvement in cell state regulation.

Next, we assessed whether these *in vitro* molecular changes are mirrored *in vivo* using the MEL006 melanoma PDX model, suitable for studying non-genetic therapeutic mechanisms^2^. Similarly, to the *in vitro* setting, short-term treatment with conoidin A, dabrafenib plus trametinib, or their combination was investigated to examine early changes in cell state switches *in vivo* (Figure 5E, S4C). Longer treatment periods, which could potentially show differences in tumor volumes, were not possible due to toxicity. We conducted single-nuclei RNA sequencing (snRNA-seq) of the four treatment groups (untreated, conoidin A, dabrafenib plus trametinib, and dabrafenib plus trametinib plus conoidin A). Harmony integration for batch effect removal followed by unsupervised clustering identified diverse cell states including NC-like, mesenchymal, melanocytic, transitional, angiogenic, and proliferative (Figure 5F). Treatment-specific cell type proportions revealed that dabrafenib plus trametinib treatment increased mesenchymal cells, which decreased upon conoidin A addition, aligning with our *in vitro* observations (Figure 5D). However, NC-like cell changes differed *in vivo*, not decreasing with dabrafenib plus trametinib plus conoidin A compared to dabrafenib plus trametinib alone. Further exploration suggested this discrepancy might stem from initial cell state differences between *in vitro* and *in vivo* contexts. Analysis of untreated *in vitro* and *in vivo* RNA-seq data using CytoTRACE^46^ revealed a higher differentiation potential in *in vitro* cells (Figure S4D), suggesting a more pronounced initial dedifferentiation state, partly explaining the observed experimental differences. Nonetheless, the reduction in mesenchymal cells with dabrafenib plus trametinib plus conoidin A treatment *in vivo* indicates conoidin A’s role in inhibiting the dedifferentiation trajectory or promoting differentiation. To further study these dynamic changes, we employed RNA velocity analysis (velocyto/scVelo)^47,48^, a computational pipeline built on a differential equation model of spliced/unspliced mRNAs that allows the inference of temporal expression dynamics. This analysis revealed that dabrafenib plus trametinib treatment significantly promoted differentiation towards mesenchymal and melanocytic states, while the transition towards a mesenchymal state was markedly reduced by addition of conoidin A, accompanying a significant decrease in mesenchymal cell numbers (Figure 5G). In summary, both *in vitro* and *in vivo* data demonstrate that MAPK inhibitor therapy induced various persisting cell states, while conoidin A co-treatment inhibited dedifferentiation and favored a shift towards a more differentiated melanocytic state, suggesting potential new therapeutic strategies and approaches.

## Discussion

The pronounced phenotypic plasticity of melanoma and resulting tumor cell state heterogeneity create multiple advantages for melanoma cells to secure their survival^11^. Previous work by us has demonstrated that some melanoma cells that persist under MAPK inhibitor therapy reside in a differentiated cell state^13^. Characteristic of this cell state, is an increased expression of the epigenetic regulator KDM5B, which triggers a shift from glycolysis-driven to OXPHOS-driven energy supply^8,16^. However, despite multiple efforts to therapeutically exploit this potential metabolic vulnerability using pharmacological OXPHOS inhibitors, improved therapy outcomes in patients have not been achieved due to rapidly emerging metabolic adaptations and the significant toxicity associated with these inhibitors^19,20^. Also, the direct inhibition of KDM5s by chemical compounds has only shown limited efficacy in terms of melanoma growth inhibition and only a minor influence on the cellular redox potential in melanoma cells^16^. This limitation is partly due to the fact that KDM5 inhibitors are typically α-ketoglutarate mimetics. As such, they compete with the KDM cofactor α-ketoglutarate, which is regulated in multiple ways, e.g. via glutamine metabolism which is often excessively utilized in tumors^49–52^. Thus, the identification of new metabolic vulnerabilities still remains a crucial strategy for advancing melanoma therapy.

Due to their high OXPHOS activity and concurrent mitochondrial electron leakage, melanoma DTPs often exhibit elevated intracellular ROS levels, which represents another potential metabolic vulnerability^8,15^. Here, we identified the PRDX/TXN ROS detoxification system as a novel metabolic target downstream of KDM5B. Under normal physiological conditions, particularly PRDXs are capable of sensing and regulating H_2_O_2_ levels locally by transferring their oxidation state to effector proteins through their sensor-transducer function^42^. In our melanoma models, inhibition of the PRDX/TXN system consequently led to increased ROS levels. Furthermore, the combination of PRDX/TXN inhibitors (auranofin, PX-12, celastrol, or conoidin A) plus MAPK-targeted melanoma drugs yielded in improved killing of cells with high KDM5B expression. As this cytotoxicity was attributed to ROS-induced cell death, it could be rescued by the addition of the ROS scavenger NAC. Similar to our study in BRAF-mutant melanoma cells, the principle of ROS-induced cell death was recently investigated also in NRAS-mutated, MEKi-resistant mouse melanoma xenografts, showing tumor growth inhibition and metastasis reduction upon application of the copper chelator neocuproine^22^.

Another type of - mainly lipid ROS-mediated - DTP cell vulnerability, termed ferroptosis, has been lately investigated in various cancer entities, including breast, ovarian, lung cancer, and melanoma^12,29,30^. Lipid ROS-inducing drugs such as the GPX4 inhibitor RSL3 are particularly effective in enhancing ferroptotic cell death in dedifferentiated, mesenchymal DTPs, but are generally limited by insufficient bioavailability^12,29,30^. Our findings now provide an important alternative targeting strategy that specifically takes into consideration melanoma cell state heterogeneity and longitudinal cell state evolution. We demonstrate that pharmacologic PRDX inhibition can induce cell death in differentiated melanoma DTPs through an alternative ROS induction pathway that is independent of ferroptosis and potentially applicable in living organisms.

Importantly, the inhibition of the KDM5B-PRDX axis by conoidin A additionally affected the cellular differentiation dynamics of melanoma DTPs under continuous dabrafenib plus trametinib (MAPK inhibitor) treatment. We observed a reduced transition of melanoma cells into the mesenchymal DTP state under concomitant conoidin A treatment both in our *in vitro* and *in vivo* transcriptomic data. In contrast, the emergence of the NC-like DTP state varied between the two experimental setups, being blocked in the *in vitro* setting, while favored in the *in vivo* setting. Reconsidering the differentiation potentials of these two melanoma models, we found that this alleged discrepancy could be ascribed to the distinct differentiation states of the models at the onset of treatment. While the melanoma cell line WM3734 resides in a dedifferentiated transcriptional state, the MEL006 PDX model is characterized by a more differentiated profile. In sum, this observation emphasizes not only the potential of PRDX-mediated ROS-targeting as a therapeutic strategy to quantitatively eliminate DTP cells, but also that ROS manipulation of tumors has to be qualitatively considered in the context of the respective cell differentiation state at therapy initiation and potential effects on cell state evolution over time. However further studies are needed now to elucidate how the inhibition of PRDXs mechanistically affects cell state shifts. For example, sensor-transducer functions of PRDXs beyond their ROS detoxification function, which are far less studied, are conceivable. These functions have been demonstrated in the past for covalent interactions with ASK1 or via PRDX1-mediated oxidation of APE1, subsequently preventing the DNA binding activity of the transcription factor nuclear factor κB (NF-κB)^42,53,54^. An essential observation of our study was that high cellular KDM5B expression fosters high PRDX/TXN gene expression. However, transcriptional activation of both KDM5B and downstream target genes appears contradictory to KDM5 demethylase activity, since H3K4 demethylation is generally associated with transcriptional repression^49^. Our ChIP-seq analysis suggests that the regulation of PRDX/TXN genes is independent of KDM5B’s demethylase activity and probably rather mediated by direct interactions with or within multi-protein epigenetic complexes containing HDAC proteins. This observation, along with other reports, supports demethylase-independent functions as a central feature of KDM5B. For example, Zhang et al. recently showed that KDM5B induces an anti-tumor immune response through the interaction with the methyltransferase SETDB1 without altering H3K4me3 levels^17^. Certainly, global epigenetic studies have to be pursued now to fully and functionally characterize KDM5B-associated chromatin complexes in melanoma.

In summary, our data suggest that targeting the PRDX/TXN ROS detoxification system could offer a promising alternative strategy downstream of KDM5B to overcome MAPK inhibitor resistance in melanoma patients. However, to exploit the full potential of pharmacological PRDX/TXN inhibition, improvements in target specificity are necessary. For example, the PRDX inhibitor mostly used in our study, conoidin A, is known to target predominantly PRDX1, PRDX2, and PRDX4. Celastrol inhibits PRDX2 and unspecific targets such as the proteasome or HSP90^55–60^. In contrast, our data suggest a strong role of PRDX3 in promoting melanoma resistance to BRAF inhibition, which is largely spared by currently available PRDX-inhibiting compounds. In addition, it should be emphasized again that melanomas consist of several coexisting DTP states; i.e. some DTP cells may not be affected by PRDX inhibition and rather be more susceptible to ferroptosis induction. Therefore, the eradication of all melanoma DTP states in the future will require both the in-depth characterization of the melanoma cell state differentiation potential at therapy onset and the individualized, probably sequential selection of different ROS-targeting strategies comprising PRDX/TXN-specific and ferroptosis inducing agents.

## STAR METHODS

### RESOURCE AVAILABILITY

#### Lead contact

Requests for further information and resources should be directed to and will be fulfilled by the lead contact Alexander Roesch.

#### Materials availability

This study did not generate new unique reagents. WM3734^KDM5Bprom-EGFP^, and the inducible Tet-On WM3734^Tet-scramble^ and WM3734^Tet-shKDM5B^ cell lines are available from the lead contact upon request via institutional material transfer agreement procedures.

#### Data and code availability

- The bulk RNA-seq data discussed in this study has been deposited in NCBI’s Gene Expression Omnibus^61^ database under accession code GSE168192. Other data reported in this paper will be shared by the lead contact upon reasonable request.
- This paper does not report original code.
- Any additional information required to reanalyze the data reported in this paper is available from the lead contact upon request.

### EXPERIMENTAL MODEL AND STUDY PARTICIPANT DETAILS

#### Melanoma cell lines and culture

Human melanoma Wistar cell line WM3734 (RRID:CVCL_6800; female sex; *BRAF^V600E^*, *PTEN^del^*, *NRAS^wt^*), and *WM3734^KDM5Bprom-EGFP ref.3^*, *WM3734^Tet-scramble^* and *WM3734^Tet-shKDM5B^* ^ref.10^ cell lines were maintained in Tu2% medium (400 ml MCDB153 basal medium, 100 ml L15 Leibovitz, 10 ml fetal bovine serum, substituted with 250 µl human insulin (10 mg/ml), 560 µl CaCl_2_ (1.5 M), and 6.8 mM L-glutamine) at 37°C and 5% CO_2_. Primary patient-derived melanoma cell lines Ma-Mel-61a (RRID:CVCL_291; male sex; *BRAF^V600E^, PTEN^wt^, NRAS^wt^*)^62^ and Ma-Mel-86c (RRID:CVCL_C7TP; female sex; *BRAF^V600E^, PTEN^wt^, NRAS^wt^*)^63^ were maintained in RPMI medium supplemented with 10% fetal bovine serum and 1% penicillin-streptomycin at 37°C and 5% CO_2_ (ethical committee approval no. 11-4715 (SCABIO), Medical Faculty, University Duisburg-Essen). Cells were trypsinized by using trypsin-EDTA (0.05%/0.02%) from PAN Biotech (Aidenbach, Germany). For Wistar cell lines trypsin neutralization solution from Lonza (Basel, Switzerland) was used. All cell lines were regularly screened for mycoplasma contamination (see Myco primer sequences in Table S1).

#### Patient-derived xenograft (PDX) model

The PDX experiment was performed in collaboration with TRACE (https://gbiomed.kuleuven.be/english/research/50488876/54502087/Trace) in accordance with local institutional guidelines and regulations. Studies involving human samples were approved by the UZ Leuven/KU Leuven Medical Ethical Committee (S63799 and S66000). Tumor tissue fragments from the cutaneous melanoma MEL006 PDX model, derived from a female patient and previously established and characterized at the TRACE Platform (UZ/KU Leuven), were implanted interscapularly in seven female, 13-14 weeks old NMRI nude mice (Rj:NMRI-Foxn1nu/nu, Janvier). Once tumors reached ∼1000 mm³, mice were randomized and treatments started with conoidin A (intraperitoneal injection of 10 mg per kg bodyweight in 5% DMSO with 95% PBS, every 3 days, 2 mice), dabrafenib plus trametinib (single daily oral gavage of 30 mg per kg body weight plus 0.3 mg per kg body weight in 10% DMSO in 90% PBS, 2 mice) or the combination of dabrafenib, trametinib plus conoidin A (2 mice). Tumor growth was measured until mice were sacrificed for tumor sample collection at the defined endpoint of the experiment for subsequent single-nuclei RNA sequencing analysis.

For conoidin A mice, the endpoint was when tumors reached the maximum permissible tumor volume. For dabrafenib plus trametinib and dabrafenib, trametinib plus conoidin A treated mice, the endpoint was when the tumor volumes were reduced by approximately 50-75% during treatment. One mouse was sacrificed before treatment start as an untreated control. Experiments involving animals were performed according to European guidelines and approved by the KU Leuven animal ethical committee (P164/2019).

The PDX model used in our experiments was originally derived from a female patient and tested in female mice. We acknowledge the importance of sex balance in experimental research. However, our standard procedure involves developing and testing xenograft models in female mice. This practice is commonly accepted in cancer research for pre-clinical testing of tumor responses to MAPK inhibitory drugs due to its reduced variability, smaller operator-induced differences, and higher reproducibility.

#### Study approval

This study complies with all relevant ethical regulations and was approved by the ethics committee of the University Hospital Essen, University of Duisburg-Essen, Medical Faculty (ethical committee approval no. 11-4715 (SCABIO)). Studies involving human samples were approved by the UZ Leuven/KU Leuven Medical Ethical Committee (S63799 and S66000) and experiments involving animals were performed according to European guidelines and approved by the KU Leuven animal ethical committee (P164/2019).

### METHOD DETAILS

#### Drugs and chemical compounds

For treatments, the following drugs and chemical compounds were used: auranofin (AdipoGen, Fuellinsdorf, Switzerland), celastrol (Selleckchem, Houston, TX, USA), conoidin A (Biomol, Hamburg, Germany), dabrafenib (Selleckchem, Houston, TX, USA), DMSO (Roth, Karlsruhe, Germany), doxycycline (AppliChem, Omaha, NE, USA), liproxstatin-1 (Cayman Chemical, Ann Arbor, MI, USA), N-acetyl-L-cysteine (NAC; Merck, Darmstadt, Germany), PLX4720 (Selleckchem, Houston, TX, USA), PX-12 (AdipoGen, Fuellinsdorf, Switzerland), RSL3 (MedChemExpress, Monmouth Junction, NJ, USA), trametinib (Selleckchem, Houston, TX, USA).

#### Western blot

For Western blot analysis whole cell lysates were prepared according to the previously described REAP protocol^64^. 15 µg of protein were separated on a 4–15% Mini-PROTEAN® TGX Stain-Free™ Protein Gel (Bio-Rad, Hercules, CA, USA) and transferred onto a PVDF membrane (Roth, Karlsruhe, Germany), following membrane blocking for 1 hour in phosphate-buffered saline (PBS) containing 0.1% Tween-20 (PBS-T) and 5% milk. Primary antibodies KDM5B (diluted 1:500, NB100-97821, Novus Biologicals, St. Louis, MO, USA and Tubulin (diluted 1:1000, 2148 Cell Signaling Technology, Cambridge, UK) were incubated in PBS-T and 5% milk at 4°C overnight. Membranes were washed with PBS-T and incubated with secondary antibody horseradish peroxidase-conjugated anti-rabbit (diluted 1:10,000, 115-035-046, Jackson Immuno Research Laboratories, West Grove, PA, USA) in PBS-T and 5% milk for 1 hour at room temperature. After washing the membranes in PBS-T, protein bands were visualized by enhanced chemiluminescence system (WesternBright Chemiluminescence Substrate, Advansta, Menlo Park, CA, USA) and captured using a FUJI LAS3000 system or Amersham™ Imager 600. Digital quantitation was performed with ImageJ 1.52i software. KDM5B signals were normalized to Tubulin signals.

#### Flow Cytometry

Cell survival and concurrent KDM5B promoter-driven EGFP signal was determined in WM3734^KDM5Bprom-EGFP ref.3^ cells by flow cytometry. For siRNA knockdown, 100,000 WM3734^KDM5Bprom-EGFP^ cells were seeded per well in a 6-well plate and allowed to attach for 24 hours. Then, siRNA knockdown was performed by transfection of siRNAs for 4 hours (Figure 3A) or 24 hours (Figure S3A/S3D/S3E), before cells were treated with either DMSO or 2.5 µM PLX4720 (Figure 3A) or 25 µM PLX4720 (Figure S3E) for 72 hours. For drug treatments without prior siRNA transfection, 50,000 WM3734^KDM5Bprom-EGFP^ cells were seeded per well in a 6-well plate and allowed to attach for 24 hours and treated with the following drugs for 72 hours: 0.1 µM celastrol, 0.5 µM conoidin A, 0.25 µM auranofin, 2.5 µM PX-12 or combined with 25 µM PLX4720 (Figure 3C), or 100 nM dabrafenib and 20 nM trametinib (Figure S3C). Subsequently, cells were harvested by trypsin-EDTA (0.05%/0.02%) and cell staining buffer (BioLegend, San Diego, CA, USA). Prior to flow cytometric measurement, 5 µl of 7-AAD (eBioscience) was added to the cells and signals were measured with Gallios flow cytometer (Beckman Coulter, Krefeld, Germany) or BD FACSAria™ III (BD Biosciences, Franklin Lakes, NJ, USA). Data analysis was performed using FlowJo_v10.8.1 software and gates were set based on the DMSO control.

To determine reactive oxygen species (ROS), 300,000 cells per 6 cm dish were seeded and allowed to attach for 24 hours. Treatments were performed for 24-72 hours with single and/or combination treatments of DMSO, 500 µM NAC, 2.5 µM conoidin A, 0.6 µM celastrol, 10 µM PX-12, 2 µM auranofin or 25 µM PLX4720. Cells were harvested and stained with 5 µM CellROX™ Green (Thermo Fisher Scientific, Waltham, MA, USA) for 30 min at 37°C. Cells were washed with PBS, resuspended in SYTOX™ Blue (Thermo Fisher Scientific, Waltham, MA, USA) in PBS (diluted 1:20.000) and analyzed by flow cytometry using BD FACSAria™ III (BD Biosciences, Franklin Lakes, NJ, USA). Data was analyzed using FlowJo_v10.8.1 and mean fluorescence signals from living cells were normalized to DMSO control samples of the respective time point.

For lipid ROS measurement, C11 BODIPY 581/591 (Cayman Chemical, Ann Arbor, MI, USA) was used. Briefly, 50,000 cells per well were seeded in a 6-well plate and allowed to attach for 24 hours. Treatments were performed for 24-72 hours with single and/or combination treatments of DMSO, 2.5 µM conoidin A, 1 µM RSL3, 10 µM liproxstation-1, and 25 µM PLX4720. During the last 30 minutes of treatment, C11 BODIPY 581/591 was added at 5 µM to each well. Then, cells were harvested and washed with PBS. Both, oxidized and non-oxidized C11 BODIPY 581/591 signals were measured at Gallios flow cytometer (Beckman Coulter, Krefeld, Germany). Data analysis was performed using FlowJo_v10.8.1 software and oxidized (FITC) to non-oxidized (PE) median fluorescence intensity (MFI) ratio was calculated for each sample. The data was normalized to DMSO control sample of the respective time point.

#### siRNA knockdown studies

Flexitube siRNAs were purchased from Qiagen (Hilden, Germany) and transfected using jetPRIME® (Polyplus, Illkirch, France) according to manufacturer’s protocol. Used siRNAs are listed in Table S2 with respective concentrations. After transfection, knockdown efficiency was confirmed by quantitative PCR (qPCR).

#### Real-time quantitative RT-PCR (qPCR)

Samples for qPCR were either prepared from siRNA-transfected WM3734, Ma-Mel-86c, Ma-Mel-61a, or WM3734^KDM5Bprom-EGFP^ cells or from longitudinally collected WM3734 samples under treatment with 0.5 µM conoidin A, 0.5 nM dabrafenib plus 0.1 nM trametinib, or 0.5 nM dabrafenib plus 0.1 nM trametinib plus 0.5 µM conoidin A. For all qPCR experiments, total RNA was isolated using the RNeasy Mini Kit according to manufacturer’s protocol (Qiagen, Hilden, Germany). 10-20 ng of RNA was used for qPCR with Luna Universal One-Step RTqPCR Kit according to manufacturer’s protocol (New England Biolabs, Ipswich, MA, USA) in a StepOnePlus Real-Time PCR system (Thermo Fisher Scientific, Waltham, MA, USA). Thermal cycler conditions were 55°C for 10 min, 95°C for 1 min, then 40 cycles of 10 sec at 95°C and 30 sec at 60°C. StepOne software v2.3 was used for data analysis. mRNA expression was calculated using the 2-DDCT method. 18S was used as housekeeping gene. All primer sequences are shown in Table S1.

#### Cell viability assessments

Short-term cell proliferation was measured using the MTT cell viability assay. 2,000 WM3734 cells were seeded in a 96-well plate and allowed to attach for 24 hours. Cells were treated with control DMSO, or different drug concentrations of 0.05 μM - 20 μM conoidin A, 0.05 μM - 20 μM celastrol, 0.01 μM - 10 μM auranofin, or 0.01 μM - 20μM PX-12 for 24, 48, and 72 hours. MTT reagent was added and metabolized MTT measured by using a spectrophotometer. Data were log transformed, normalized to DMSO control of each time point and depicted as nonlinear regression.

For 2D clonogenic assay 27,000 Ma-Mel-86c cells were seeded in a 24-well plate and allowed to attach for 24 hours. Drug treatment with control DMSO, 500 µM NAC, 25 µM PLX4720, 2.5 µM conoidin A and combinations of the mentioned drugs was performed for 72 hours. Then cells were fixed and stained with crystal violet before pictures were taken. After dissolving with 70% ethanol, crystal violet absorption was measured at OD_550nm_.

Long-term proliferation assays were performed in a Nyone® cell imager (Synentec, Elmshorn, Germany). Briefly, 10,000 cells per well were seeded in a 24-well plate and allowed to attach for 24 hours. Ma-Mel-61a cells were continuously treated with 0.5 µM conoidin A or DMSO until cells grew confluent, while treatment with 1 µM PLX4720 and the combination of 1 µM PLX4720 and 0.5 µM conoidin A was continued for 38 days. WM3734 cells were continuously treated with control DMSO, 0.5 µM conoidin A, 0.2 µM PLX4720 or the combination of 0.2 µM PLX4720 and 0.5 µM conoidin A for 51 days or with DMSO, 0.5 µM conoidin A, 1 nM dabrafenib and 0.2 nM trametinib or the triple combination of 1 nM dabrafenib and 0.2 nM trametinib and 0.5 µM conoidin A for 14 days. During the experiments, medium was refreshed every 2-3 days and confluence was recorded in the Nyone® cell imager. On the last day of the experiment, cells were stained in the well with Hoechst (10 ml/mg diluted 1:2,000 in medium, Thermo Fisher Scientific, Waltham, MA, USA) for 30 min at 37°C, washed and counted with Nyone® cell imager. Confluence measurements and nuclei counts were visualized using GraphPad Prism version 8.4.3.

#### Chromatin immunoprecipitation sequencing (ChIP-seq)

H3K4me3-ChIP-seq was performed according to a previously described protocol^16^. Briefly, WM3734^Tet-shKDM5B^ knockdown and WM3734^Tet-scramble^ control cells were used and knockdown was induced with 500 ng/ml doxycycline for 72 hours. Cells were harvested, and aliquots of 2 × 10^6^ cells were fixed with 1% formaldehyde for 6 min at room temperature, followed by 125 mM glycine for 5 min. Cells were washed in ice-cold PBS, resuspended, centrifuged, and cell pellets were snap-frozen. After cell lysis, chromatin was sheared using an ultrasonicator with 50W power, 20% duty factor, and 200 cycles per burst for 3:30 min. For immunoprecipitation, 1 µl H3K4me3 (Abcam) or 1 µl IgG (Dianova) antibody was coupled to magnetic Dynabeads protein A and G (each 5.5 µl) for 2 hours at 4°C and 40 rpm. Sheared chromatin from 1 × 10^6^ cells was added to the antibody-coupled beads and incubated overnight at 4°C and 40 rpm. Beads were washed in RIPA buffer, and crosslinking reversed by RNase digestion and proteinase K treatment. DNA was purified using the QIAquick PCR purification Kit (Qiagen, Hilden, Germany). ChIP-seq libraries were prepared from 5 ng of ChIP DNA using the Next Gen DNA Library Kit (Active Motif, Carlsbad, CA, USA) according to manufacturer’s protocol. The libraries were quantified with a Qubit 4 fluorometer (Invitrogen) and fragment length was characterized on a TapeStation D1000 (Agilent, Santa Clara, CA, USA). Samples were sequenced on an Illumina NextSeq500 with single-end 85bp reads. Fastq files were quality checked and trimmed using the FastQC and Cutadapt algorithms as implemented in trigalore. Cleaned reads were mapped to the human genome (GRCh38 build) using bowtie2 with sam output format. The hg38 human genome parameter was set into the Homer environment and peaks were then called using Homer with style set to “histone”. Peak annotation motif identification and differential binding were all performed in Homer and bedgraph files were generated for visualization.

KDM5B-ChIP-seq was performed in collaboration with Active Motif (Carlsbad, CA, USA). Approximately 3 × 10^7^ Ma-Mel-61a or 1.2 × 10^8^ Ma-Mel-86c cells were prepared according to the ChIP Cell Fixation Protocol from Active Motif. Briefly, cells were fixed in 1% formaldehyde solution for 15 min at room temperature, followed by the addition of glycine solution for 5 min at room temperature to stop the fixation. Cells were scraped from the culture surface, washed once in PBS-Igepal, and once in PBS-Igepal containing 1 mM PMSF. Active Motif prepared chromatin, performed ChIP reactions, generated libraries, sequenced the libraries and performed basic data analysis. Briefly, chromatin was isolated by adding lysis buffer and using a dounce homogenizer, following sonication and DNA shearing to an average length of 300-500 bp using Active Motif’s EpiShear probe sonicator (#53051, Active Motif, Carlsbad, Ca, USA). Genomic DNA (Input) was prepared by treating aliquots of chromatin with RNase, proteinase K and heat for de-crosslinking, followed by SPRI beads clean up (Beckman Coulter) and quantitation by Clariostar (BMG Labtech). 30-40 µg of chromatin was precleared with protein G agarose beads (Invitrogen) and genomic DNA regions of interest were isolated using 15 µl of KDM5B antibody (either Cell Signaling Technology (CST, #15327), or Sigma-Aldrich (Sigma, #HPA027179), or a mixture of both). Complexes were washed, eluted from the beads with SDS buffer, and subjected to RNase and proteinase K treatment. Crosslinks were reversed by incubation overnight at 65°C, and ChIP DNA was purified by phenol-chloroform extraction and ethanol precipitation. ChIP DNAs were processed into Illumina sequencing libraries, quantified, and sequenced on Illumina’s NovaSeq 6000 (75 nt reads, single-end). Reads were aligned to the human genome (hg38) using the BWA algorithm (default settings). Duplicate reads were removed and only uniquely mapped reads (mapping quality >= 25) were used for further analysis. Alignments were extended *in silico* at their 3’-ends to a length of 200 bp, which is the average genomic fragment length in the size-selected library, and assigned to 32-nt bins along the genome. The resulting histograms (genomic “signal maps”) were stored in bigWig files. Peak locations were determined using the MACS algorithm (v2.1.0) with a cutoff of *p*-value = 1e-7. Peaks that were on the ENCODE blacklist of known false ChIP-seq peaks were removed.

#### Chromatin immunoprecipitation and quantitative PCR (ChIP-qPCR)

For ChIP-qPCR, the SimpleChIP® Enzymatic Chromatin IP Kit (Cell Signaling Technology, Cambridge, UK) was used according to manufacturer’s protocol with minor modifications such as an additional cross-linking step with 1.5 mM EGS in cold PBS for 20 min at room temperature prior to formaldehyde cross-linking. Chromatin digestion and nuclei preparation were performed using an optimized volume of 0.25 µl micrococcal nuclease enzyme per ChIP for 10 min at 37°C. ChIP was performed with 3 µg of IgG (negative control) or KDM5B (#15327, Cell Signaling Technology) antibodies from 10 µg chromatin. Primers were designed within KDM5B binding areas and used for analysis of purified DNA using qPCR analysis (see Table S1). The % Input method was used for normalization.

#### Sample preparation for immunoprecipitation mass spectrometry

For immunoprecipitation mass spectrometry, nuclear cell lysates from WM3734 melanoma cells were generated by using nuclear complex Co-IP Kit (#54001, Active Motif, Carlsbad, CA, USA) according to the manufacturer’s protocol. Briefly, cells were harvested using a cell scraper. Cell lysis was performed in hypertonic buffer by adding detergent. Nuclei were lysed for 90 min at 4°C and protein/protein interactions were preserved in low-salt buffer. 500 μg of the nuclear extract were used per IP reaction and mixed with either control IgG (Dianova) or anti-KDM5B (#15327, Cell Signaling technology) antibodies for 2 hours at 4°C rotating at 40 rpm, subsequently, magnetic Dynabeads protein A were added and incubated for 2 hours at 4°C rotating at 40 rpm. Beads were washed twice in IP wash buffer for 4 min at 4°C rotating at 40 rpm and eluted in two steps using 50 μl of 2x concentrated and 1x concentrated elution buffer (4x SP3 lysis and elution buffer: 4% (w/v) SDS, 40 mM TCEP, 160 mM Chloroacetamide, 200 mM HEPES pH 8) for 5 min at 95°C). 10 µl aliquots were used to confirm KDM5B protein precipitation in Western blot analysis using KDM5B antibody (#NB100-97821, Novus Biologicals, St. Louis, MO, USA) and clean-blot reagent (#21230, Thermo Fisher Scientific, Waltham, MA, USA). The remaining 90 µl were used for further processing.

#### Single-pot solid-phase-enhanced sample preparation (SP3) for LC/MS

Sample preparation for LC/MS/MS is based on the SP3 protocol^65^. The eluted proteins (90 µl) were cooled down to room temperature and then supplemented with 150 µg hydrophobic (#65152105050250) and 150 µg hydrophilic (#45152105050250) SeraMag Speed Beads (Cytiva) and gently mixed for 1 min. Then 100 µl 100% vol/vol Ethanol (EtOH) was added before incubation for 20 min at 24°C shaking vigorously. The beads were collected on a magnet and the supernatant aspirated. The beads were then washed 4 times with 180 µl 80% EtOH (collection time on the magnet minimum of 4 min). The beads were finally taken up in 100 µl 25 mM ammoniumbicarbonate (ABC) containing 1 µg Trypsin (Protein:Trypsin ratio 30:1). To help bead dissociation, samples were incubated for 5 min in a sonification bath (preheated to 37°C). Samples were incubated overnight, shaking vigorously (1,300 rpm). Next day samples were acidified with formic acid (FA, final 1% vol/vol) before collection on a magnet. The supernatants were transferred to a fresh Eppendorf tube, before removing trace beads using a magnet for 5 min. The tryptic digests were then desalted on home-made C18 StageTips as described^66^. Briefly, peptides were immobilized and washed on a 2 disc C18 StageTip. Samples were then dried using a vacuum concentrator (Eppendorf, Hamburg, Germany) and the peptides were taken up in 0.1% formic acid solution (15 μl) and directly used for LC-MS/MS experiment.

#### LC-MS/MS

Experiment was performed on an Orbitrap Fusion Lumos (Thermo Fisher Scientific, Waltham, MA, USA) that was coupled to an EASY-nLC 1200 liquid chromatography (LC) system (Thermo Fisher Scientific, Waltham, MA, USA). The LC was operated in the one-column mode. The analytical column was a fused silica capillary (75 µm × 28 cm) with an integrated fused silica capillary with an integrated sintered frit (FossilIontech) packed in-house with Kinetex C18-XB core shell 1.7 µm resin (Phenomenex). The analytical column was encased by a column oven (Sonation) and attached to a nanospray flex ion source (Thermo Fisher Scientific, Waltham, MA, USA). The column oven temperature was adjusted to 50°C during data acquisition. The LC was equipped with two mobile phases: solvent A (0.2% formic acid, FA; 97.8% H_2_O; 2% Acetonitrile, ACN) and solvent B (0.2% FA; 80% ACN; 19.8% H_2_O). All solvents were of UPLC grade (Honeywell). Peptides were directly loaded onto the analytical column with a maximum flow rate that would not exceed the set pressure limit of 980 bar (usually around 0.5 – 0.7 µl/min). Peptides were subsequently separated on the analytical column by running a 105 min gradient of solvent A and solvent B (start with 3% B; gradient 3% to 9% B for 6:30 min; gradient 9% to 30% B for 62:30 min; gradient 30% to 50% B for 24 min; gradient 50% to 100% B for 2:30 min, and 100% B for 9:30 min) at a flow rate of 350 nl/min. The mass spectrometer was operated using Tune v3.3.2782.28. The mass spectrometer was set in the positive ion mode. Precursor ion scanning was performed in the Orbitrap analyzer (FTMS; Fourier Transform Mass Spectrometry) in the scan range of m/z 375-1500 and at a resolution of 240000 with the internal lock mass option turned on (lock mass was 445.120025 m/z, polysiloxane)^67^. Product ion spectra were recorded in a data dependent fashion in the ITMS at “rapid” scan rate. The ionization potential (spray voltage) was set to 2.5 kV. Peptides were analyzed using a repeating cycle consisting of a full precursor ion scan (AGC standard; max acquisition time “Auto”) followed by a variable number of product ion scans (AGC 300% and acquisition time auto) where peptides are isolated based on their intensity in the full survey scan (threshold of 4000 counts) for tandem mass spectrum (MS2) generation that permits peptide sequencing and identification. Cycle time between MS1 scans was 3 sec. Fragmentation was achieved by stepped Higher Energy Collision Dissociation (sHCD) (NCE 27, 32, 40). During MS2 data acquisition dynamic ion exclusion was set to 20 sec and a repeat count of one. Ion injection time prediction, preview mode for the FTMS, monoisotopic precursor selection and charge state screening were enabled. Only charge states between +2 and +7 were considered for fragmentation.

#### Peptide and Protein Identification using MaxQuant

RAW spectra were submitted to an Andromeda^68^ search in MaxQuant (2.0.3.0.) using the default settings^69^. Label-free quantification and match-between-runs was activated^70^. The MS/MS spectra data were searched against the Uniprot *A. thalian*a reference proteome (UP000005640_2-HomoSapiens(08-2022).fasta; 79759 entries). All searches included a contaminants database search (as implemented in MaxQuant, 245 entries). The contaminants database contains known MS contaminants and was included to estimate the level of contamination. Andromeda searches allowed oxidation of methionine residues (16 Da) and acetylation of the protein N-terminus (42 Da). Carbamidomethylation on Cystein (57 Da) was selected as static modification. Enzyme specificity was set to “Trypsin/P”. The instrument type in Andromeda searches was set to Orbitrap and the precursor mass tolerance was set to ±20 ppm (first search) and ±4.5 ppm (main search). The MS/MS match tolerance was set to ±0.5 Da. The peptide spectrum match FDR and the protein FDR were set to 0.01 (based on target-decoy approach). For protein quantification unique and razor peptides were allowed. Modified peptides were allowed for quantification. The minimum score for modified peptides was 40. Label-free protein quantification was switched on, and unique and razor peptides were considered for quantification with a minimum ratio count of 2. Retention times were recalibrated based on the built-in nonlinear time-rescaling algorithm. MS/MS identifications were transferred between LC-MS/MS runs with the “match between runs” option in which the maximal match time window was set to 0.7 min and the alignment time window set to 20 min. The quantification is based on the “value at maximum” of the extracted ion current. At least two quantitation events were required for a quantifiable protein. Further analysis and filtering of the results was done in Perseus v1.6.10.0.^71^. Comparison of protein group quantities (relative quantification) between different MS runs is based solely on the LFQ’s as calculated by MaxQuant, MaxLFQ algorithm^70^.

#### Bulk RNA-seq transcriptional profiling

For generating total RNA, knockdown WM3734^Tet-shKDM5B^ and WM3734^Tet-scramble^ control cells (175,000 cells) were seeded in a 6 cm dish and induced with 500 ng/ml doxycycline over 72 hours. Total RNA was isolated using RNeasy Mini Kit according to the manufacturer’s protocol (Qiagen, Hilden, Germany). Barcoded stranded mRNA-seq libraries were prepared using NEB Ultra II stranded RNA library prep kit from 1 µg of purified total RNA according to the manufacturer’s protocol (New England Biolabs, Ipswich, MA, USA). Libraries were loaded on the Illumina sequencer NextSeq 500 and sequenced uni-directionally, generating ∼550 million reads 85 bases long. Fastq files were quality checked with the Fastqc tool and the reads were trimmed with Trimmomatic tool. Reads were mapped to the human genome (GRCh38 build) using STAR with the quantmode activated for feature quantification. Gene expression matrices with raw counts were processed with Deseq2 for normalization and differential gene expression.

#### Sample preparation for *in vitro* single-cell RNA sequencing (scRNA-seq)

For scRNA-sequencing, WM3734 cells were seeded and allowed to attach for 24 hours. Then, cells were either harvested (untreated day 0 sample) or treated with 0.5 µM conoidin A, 0.5 nM dabrafenib plus 0.1 nM trametinib or the triple combination of 0.5 µM conoidin A, 0.5 nM dabrafenib, and 0.1 nM trametinib for 4 days. Sample aliquots were stored in freezing medium in liquid nitrogen until further processing. Approximately 8,000 cells per sample were loaded onto Chromium Controller (10x Genomics, Pleasanton, CA, USA). scRNA-seq libraries were generated using the Chromium Next GEM Single Cell 3’ Kit v3.1 kit (10x Genomics, Pleasanton, CA, USA), and Next GEM Chip G Single Cell kit (10x Genomics, Pleasanton, CA, USA) according to the manufacturer’s protocol. cDNA traces were amplified with 12 cycles and GEX libraries were constructed with the Dual Index Kit TT Set A according to manufacturer’s protocol. A total of 5,000 cells per samples were targeted and an Agilent 2100 Bioanalyzer with a high-sensitivity DNA kit (Agilent, Santa Clara, CA, USA) was used to assess the quality of cDNA traces and final gene expression libraries. The resulting libraries were pooled together according to balanced amounts and paired-end sequencing was performed on a NextSeq2000 with 50,000 read pairs per cell.

#### Sample preparation for *in vivo* single-nuclei RNA sequencing (snRNA-seq)

Snap-frozen tumor samples were embedded in Tissue-Tek® O.C.T and cut in slices. For RIN quality analysis, RNA was isolated with RNeasy Plus Micro Kit (Qiagen, Hilden, Germany) from 5 µm slices according to manufacturer’s protocol. Depending on the tumor size, several 20 µm slices from each tumor were used to isolate nuclei for library preparation. Isolation was performed according to previously published protocol^72^ using a dounce homogenizer with 10 times pestle A and 10 times pestle B. Nuclei were washed in total 4 times and resuspended in storage buffer before counting on LunaFX7 (BioCat, Heidelberg, Germany). Approximately 11,500 nuclei per sample were loaded onto Chromium X (10x Genomics, Pleasanton, CA, USA). SnRNA-seq libraries were generated using the Chromium Next GEM Single Cell 3’ Kit v3.1 kit (10x Genomics, Pleasanton, CA, USA) and Next GEM Chip G Single Cell kit (10x Genomics, Pleasanton, CA, USA) according to the manufacturer’s protocol. cDNA traces were amplified with 13-14 cycles and GEX libraries were constructed with the Dual Index Kit TT Set A according to manufacturer’s protocol. A total of 7,000 nuclei per samples were targeted and quality of cDNA traces and final gene expression libraries were analyzed with TapeStation and a high-sensitivity D5000 ScreenTape (Agilent, Santa Clara, CA, USA). Samples were sequenced on DNBseq platform DNBSEQ-G400 with read length PE100bp.

#### Single-cell/single-nuclei RNA sequencing data processing

Raw sequencing data (FASTQ files) were processed using the CellRanger software (version 7.0.1) with the GRCh38-2020A or mm10-2020A reference genome for mapping. The resulting gene count matrices were imported into R for subsequent analysis using the Seurat package^73^.

#### *In vitro* data analysis

For *in vitro* studies, data from four samples were integrated. Cells were filtered to retain only those meeting quality control criteria: a gene count (nFeature) range of 1,000 to 7,500 and mitochondrial gene content below 25%. This filtration yielded a total of 19,408 high-quality cells. Normalization was performed using Seurat’s SCTransform function, followed by principal component analysis (PCA) with RunPCA. Dimensionality reduction was achieved using RunUMAP, with PCA dimensions set to 1:10.

#### *In vivo* patient-derived xenograft (PDX) data analysis

The PDX samples, containing a mix of human and mouse cells, were initially mapped to both human and mouse reference genomes. Cell origin (human or mouse) was determined based on the number of human and mouse genes (nGene) per cell. Potential doublets were identified and removed using scDblFinder^74^. The data from seven samples containing human-origin cells (tumor cells) were merged. Cells were selected based on quality with criteria set at an nFeature range of 1,250 to 7,500 and mitochondrial gene content below 5%, resulting in 24,386 cells. Data normalization was conducted using the NormalizeData function in Seurat, followed by ScaleData and RunPCA. Batch effects were corrected using the Harmony algorithm. Cell clustering was performed using the FindNeighbors and FindClusters functions in Seurat (dimensions = 1:10, resolution = 0.3), and dimensionality reduction was again achieved using RunUMAP. Cell annotations were based on established literature^12,44^.

#### Data visualization and analysis

For visualization purposes, including violin plots, cell proportion bar plots, and heatmaps, the SeuratExtend Package (https://github.com/huayc09/SeuratExtend) was utilized^75^. The transcription factor network was inferred using the SCENIC workflow implemented in the Nextflow pipeline^45^. Regulon activity in each cell was quantified using the AUCell score within the Bioconductor AUCell package. Gene Set Enrichment Analysis (GSEA) was conducted using the AUCell package, focusing on gene sets referenced in previous publications^12,44^. Statistical analyses were performed using the Wilcoxon test, with *p*-value adjustments made via the Holm method.

#### Inference of differentiation trajectories

To infer temporal dynamics in scRNA-seq data, the velocyto pipeline was utilized^47^. Spliced and unspliced RNA matrices were computed from *.bam files and barcode metadata generated by CellRanger. Differentiation trajectories were visualized using UMAP embeddings, created with Seurat in R. The trajectories, depicted through arrows, were illustrated using scVelo in Python^48^.

#### In silico analysis

Correlation analysis: Gene expression from publicly available single-cell RNA-sequenced human melanomas (n = 1252 melanoma cells from 19 tumors^33^) were used for correlation analysis. Malignant cells were grouped according to KDM5B expression in high (expression value above median) and absent/low (expression value 0 plus expression values below median). Gene signatures (HALLMARK OXIDATIVE PHOSPHORYLATION, REACTOME DETOXIFICATION OF REACTIVE OXYGEN SPECIES, KEGG CITRATE CYCLE TCA CYCLE and REACTOME GLUTATHIONE CONJUGATION) were downloaded from https://www.gsea-msigdb.org/gsea/msigdb. Gene signature scores were calculated using the following equation: 1/x * SUM(expression value 1-x genes)^76^.

Kaplan-Meier overall survival analysis: Gene expression data from the TCGA_SKCM was downloaded from the UCSC Xena database^77^. Patients were grouped into low, median, or high expression by using X-Tile version 3.6.1 cut-point optimization^78^. If patients provided multiple tissue samples, only one sample per patient was utilized. Kaplan–Meier survival curves, hazard ratios (HR) with 95% confidence intervals, and *p*-values from log-rank tests were generated using GraphPad Prism version 8.4.3.

*In silico* analysis of ChIP-seq data: Publicly available KDM5B, HDAC1, and HDAC2 binding data from the ChIP-Atlas^35^ was downloaded and visualized using the Integrative Genomics Viewer version 2.11.9.

### QUANTIFICATION AND STATISTICAL ANALYSIS

Statistical significance was tested using GraphPad Prism version 8.4.3. The following *p*-values were considered statistically significant (* *p*<0.05, ** *p*<0.01, *** *p*<0.001, and **** *p*<0.0001).

Exact statistical analysis methods are described in the respective figure legends. Data are presented as mean ± SD as indicated in the figure legends. Boxplots in Figure 5 and Figure S4 were created using the ggplot2 package in R. They visualize five summary statistics (the median, two hinges, and two whiskers). The lower and upper hinges correspond to the first and third quartiles (the 25th and 75th percentiles). The upper whisker extends from the hinge to the largest value no further than 1.5 * IQR from the hinge (where IQR is the inter-quartile range, or the distance between the first and third quartiles). The lower whisker extends from the hinge to the smallest value at most 1.5 * IQR of the hinge.

## Acknowledgments

We thank M. Herlyn for providing the WM3734 cell line. We thank A. Squire and A. Brenzel from the Imaging Center Essen (IMCES), the Central Animal Facilities of the Medical Faculty Essen, V. Benes from the Genomics Core Facility Gene Core at the EMBL Heidelberg, and L. Otte, P. Braß, and the late S. Kumar for their technical support. We would also like to express our gratitude to Active Motif, particularly to Matthias Spiller-Becker, for their invaluable assistance with ChIP-seq experiments and data analysis. We would like to than E. Demesmaeker, F. Vervloesem and F. Stinkens for technical support in the PDX experiment. Servier Medical Art was used to created image in Figure 1C. Servier Medical Art by Servier is licensed under a Creative Commons Attribution 3.0 Unported License (https://creativecommons.org/licenses/by/3.0/). Biorender was used to create images in Figure 5B and Figure 5E (https://www.biorender.com/).

F.R. is funded by Melanoma Research Alliance and the Wolfgang & Gertrud Boettcher Foundation. H.C. was supported by a “Welcome Back” grant of the Medical Faculty of the UDE. This work was partly funded by the Brigitte und Dr. Konstanze Wegener-Stiftung (R.V.), BMBF DKTK ED03 (J.C.B.), EFKS UMESciA (N.S.), and the German Research Foundation (DFG, Deutsche Forschungsgemeinschaft) Project-ID 418179183 (KFO 337): RO 3577/7-1, RO 3577/7-2 (A.R.), SCHA 422/17-1, SCHA 422/17-2 (D.S.), SI 1549/3-1, SI 1549/3-2 (J.S.), BE 1394/12-1, BE 1394/12-2 (J.C.B.), PA 2376/1-1, PA 2376/1-2 (A.P.), Projekt-ID 424228829 (CRC1430, A.R., A.P., S.P., F.K.), FOR5427 SP4 (D.R.E.); EN984/15-1, 16-1 and 18-1 (D.R.E.); TR296 P09 (D.R.E.); TR332 A3 and Z1 (D.R.E.), and INST 20876/486-1 (D.R.E.).

## Author contributions

Conceptualization: S.E., H.C., F.R., and A.R.

Methodology: S.E., H.C., R.V., Y.H., S.S.L., S.M., S.S., B.S., E.L., M.F.B., J.C.M., A.P., D.R.E., L.M.B., F.M., J.K., B.T.G., S.P., S.N., F.K., J.C.B., A.T., F.R., and A.R.

Investigation: S.E., H.C., R.V., Y.H., S.S.L., S.M., N.S., V.S., S.Sch., A.H., M.F.B., F.K., and F.R.

Formal analysis: R.V., Y.H., S.S.L., J.F., and F.R.

Resources: H.C., R.V., Y.H., S.S.L., S.M., N.S., V.U., S.S., B.S., E.L., J.S., M.F.B., J.C.M., A.P., B.Sch., D.S., J.C.B., A.T., F.R., and A.R.

Project administration: F.R. and A.R.

Supervision: A.R.

Visualization: S.E. and Y.H.

Writing (original draft): S.E., H.C., and A.R.

Writing (review and editing): all authors

## Supplemental information

Figures S1-S4 and Tables S1-S2

